# Dynamic Monitoring of Antibody Drug Conjugates Targeting TROP2 or HER2 in Breast Cancer using Circulating Tumor Cells

**DOI:** 10.1101/2025.04.02.646822

**Authors:** Avanish Mishra, Rachel O. Abelman, Quinn Cunneely, Victor Putaturo, Akansha A. Deshpande, Remy Bell, Elizabeth M. Seider, Katherine H. Xu, Mythreayi Shan, Justin Kelly, Shih-Bo Huang, Olivia Rieur, Justine Knape, Kaustav A. Gopinathan, Kruthika Kikkeri, Jon F. Edd, John Walsh, Charles S. Dai, Leif W. Ellisen, David T. Ting, Linda Nieman, Mehmet Toner, Aditya Bardia, Daniel A. Haber, Shyamala Maheswaran

**Affiliations:** Center for Engineering in Medicine and Surgery, Massachusetts General Hospital and Harvard Medical School, Charlestown, Massachusetts 02129, USA; Krantz Family Center for Cancer Research, Massachusetts General Hospital Cancer Center and Harvard Medical School, Charlestown, Massachusetts, 02129, USA; Division of Hematology Oncology, Massachusetts General Hospital Cancer Center and Department of Medicine, Harvard Medical School, Boston, Massachusetts 02114, USA; Shriners Children’s Hospital, Boston, Massachusetts, 02114, USA; Howard Hughes Medical Institute, Chevy Chase, Maryland, 20815, USA

## Abstract

Antibody-drug conjugates (ADCs) target surface proteins on cancer cells, leading to internalization and delivery of a drug payload, thereby enhancing selectivity and minimizing toxicity. ADCs against TROP2 (Sacituzumab govitecan) or HER2 (T-DXd) have demonstrated efficacy in metastatic breast cancer, yet paradoxically, outside of *HER2*-amplified breast cancers, expression levels of these breast cancer-enriched epitopes in tumor biopsies have not been strongly correlated with clinical response. We undertook serial quantitative imaging of circulating tumor cells (CTCs) in a prospective cohort of 35 patients treated with either of these ADCs. At the single-cell level, expression of TROP2 and HER2 within individual patients is highly heterogeneous in both CTCs and paired tumor biopsies. Measurement of these epitopes on CTCs immediately prior to ADC therapy does not predict depth of clinical response. However, absence of CTCs or >80% reduction in CTC numbers after three weeks of treatment (CTC^Low^) predicts durable response, compared with CTC^High^ cases (TROP2: HR 5.15, P = 0.012; HER2: HR 6.01, P<0.001). Targeted epitopes are not commonly downregulated on CTCs at the time of acquired clinical resistance, and switching between TROP2- and HER2-targeting ADCs sharing similar payloads infrequently leads to second-line response. Thus, while CTC burden is correlated with response to these ADCs, the level of TROP2 or HER2 expression is poorly predictive. These findings point to sensitivity to the drug payload as a potential driver of clinical response to currently approved ADCs in breast cancer.

**SIGNIFICANCE STATEMENT:** The clinical efficacy of ADCs may depend both on differential targeting of cancer cells, using antibodies to tumor-enriched epitopes, and on the cleavage and release of drug payloads to which cancer cells are sensitive. ADCs currently used to treat breast cancer target either of two epitopes, TROP2 or HER2, but they share chemically related payloads. While CTC numbers track with response, we find that epitope expression is not strongly predictive. The limited success of sequentially switching between TROP2- or HER2-targeting ADCs as second-line treatment, following progression on a first-line ADC, highlights the need to incorporate non-cross-resistant drug payloads on ADCs to overcome such acquired resistance.

## INTRODUCTION

ADCs combine an antibody specifically targeting a tumor cell-enriched epitope with a cytotoxic payload, thereby providing the potential to deploy more selective chemotherapy with a favorable therapeutic window and toxicity profile (1). This delivery of payload to cells containing tumor-associated antigens allows the use of drugs substantially more toxic than traditional chemotherapy (2). ADCs were first deployed in breast cancer for the treatment of metastatic breast cancer (MBC) with *HER2* gene amplification (HER2+) (3). Subsequently, ADCs have been FDA-approved for advanced MBCs lacking *HER2* gene amplification, including two that target another cell surface epitope, TROP2 (sacituzumab govitecan [SG] and datopotamab deruxtecan [Dato-DXd]), and one targeting HER2 (trastuzumab deruxtecan [T-DXd]) (4–7). The promising first-line application of ADCs in combination with either immune checkpoint blockade (ASCENT-04 trial: SG + pembrolizumab vs chemotherapy + pembrolizumab for metastatic TNBC) (8) or with antibody disrupting HER2 dimerization (DESTINY-Breast09 trial: T-DXd + pertuzumab vs paclitaxel + trastuzumab + pertuzumab for HER2+ breast cancer) (9, 10) is poised to revolutionize the treatment of MBC, potentially replacing traditional chemotherapy as first-line regimens. In addition to the treatment of metastatic cancer, ADCs are being tested in the potentially curative setting of adjuvant therapy for localized breast cancer (11–13). Given the success of these ADCs, there are many more at various stages of development for breast cancer, as well as other solid tumors (7, 14, 15) and there is an urgent need to identify reliable biomarkers to help guide their selection and understand mechanisms of resistance following first-line ADC therapy.

Efforts to use tissue-based epitope expression to guide ADC selection have produced mixed results. TROP2, the target of SG and Dato-DXd, is highly expressed in normal, developing epithelial tissues compared with adult organs, and it is aberrantly expressed in certain tumors, including both triple-negative breast cancer (TNBC) and hormone receptor (HR)-positive breast cancer (16–19). Biomarker studies in tumor biopsy specimens have suggested a poor correlation between clinical response to SG and the degree of TROP2 expression at the time of initial disease presentation or subsequent metastatic relapse (13, 20, 21). A “bystander effect,” whereby ADCs recruited to TROP2-expressing cells may then kill surrounding cells lacking the antigen, has been postulated (22, 23). For ADCs targeting HER2, such as T-DXd, efficacy has also been demonstrated in HER2-expressing cancers lacking *HER2* gene amplification and its associated biological dependence on HER2 signaling. Moreover, this efficacy was subsequently shown to extend to HER2-low breast cancers (7), and even breast cancers with virtually no detectable HER2 expression (“ultra-low”) were shown to be responsive to HER2-ADCs, leading to the expansion of FDA approval, while also raising questions about the tumor epitope dependence of HER2 ADC therapy (23, 24).

While ADCs have proven to be highly effective in many patients with advanced breast cancer, other patients exhibit *de novo* resistance or acquire drug resistance after only a few months of therapy. The relative contributions of altered epitope binding versus acquired resistance to the drug payload remain to be defined (25, 26). This is particularly critical in patients who are treated sequentially with different ADCs, in whom non-cross-resistant ADCs must be deployed. Retrospective institutional series have suggested that patients with MBC who receive a first-line ADC often experience shorter treatment times with second-line ADCs (27–30). Notably, the three currently FDA-approved ADCs for breast cancer contain payloads with highly potent, albeit functionally similar, topoisomerase-I (TOP1) inhibitors (29), and acquired TOP1 gene mutations contribute to approximately 13% cases of resistance (25).

Biopsy of a metastatic lesion typically provides clinically informative data regarding the molecular characteristics of cancer in patients with highly pre-treated metastatic disease. However, sampling of a single metastatic lesion, selected on the basis of its accessibility to biopsy, may not fully represent the spatial and temporal heterogeneity of epitope expression across multiple tumor deposits that are present in patients with metastatic breast cancer. Progressive lesions may be inaccessible, and accessible bone metastases often yield poor-quality specimens due to the need for decalcification. In this setting, non-invasive liquid biopsies analyzing CTCs may address multiple challenges, enabling capture of cancer cells in the blood representing multiple sites of disease, as well as allowing non-invasive serial analysis of protein expression from patients receiving ADC therapies (31–34).

Multiple technologies have been employed for CTC isolation from blood samples, and their underlying cancer cell enrichment principles may influence the measurement of cell surface epitopes. For instance, capture of CTCs based on their expression of the epithelial cell surface markers EpCAM may lead to selection bias for co-expressed epithelial markers, such as TROP2 or HER2. Alternatively, epitope-independent size-based isolation of CTCs may fail to detect many CTCs whose cell diameter overlaps with that of leukocytes (34). To test the relationship between TROP2 and HER2 protein expression on cancer cells and clinical response to their targeting ADCs, we used a microfluidic technology that effectively depletes red blood cells and platelets, and then removes leukocytes using magnetically conjugated antibodies, thus enriching CTCs without bias for either size or epithelial differentiation (35–39). Among CTCs isolated, the application of multispectral fluorescent imaging for the ADC target epitopes of interest enables single-cell analysis of tumor cell heterogeneity. We applied this technology for serial monitoring in 35 patients with breast cancer (38 treatment courses) during treatment with either TROP2 or HER2 targeting ADCs, or sequentially with the two ADCs.

## RESULTS

### Quantitative imaging of TROP2 and HER2 epitope expression in CTCs using multispectral imaging

We have previously described a microfluidic leukocyte-depletion platform (CTC-iChip) that first excludes non-nucleated cells (RBCs and platelets) and then achieves efficient removal of magnetically tagged WBCs using antibodies against the common leukocyte epitopes CD45, CD16, and CD66b, thereby providing unbiased enrichment for CTCs (34, 35, 37, 38) (**Figure 1A**). The enriched CTCs themselves are free of conjugated antibodies and magnetic beads, providing optimal resolution for quantitative imaging. While the CTC-iChip platform previously required the immediate processing of freshly collected blood specimens, we adapted the analytical technology to allow for the fixation of blood specimens at the time of collection, thereby increasing the time available for specimen processing to up to 72 hours (**Supplementary Figure 1**). This was accomplished by collecting blood in Streck tubes that enable uniform light fixation and by imaging cells using a multispectral microscopy platform with subcellular resolution (PhenoImagerHT, Akoya Biosciences). Applying this revised processing approach, we also observed that reducing the concentration of leukocyte-tagging antibodies and magnetic particles by up to fourfold results in high CTC enrichment, with only a modest reduction in CTC purity, without compromising single-cell CTC imaging and enumeration (**Supplementary Figure 1B**). Modeling spiking of cultured CTCs into Streck-fixed blood samples stored for 24 hours shows a CTC recovery of 96.9 ± 5.1%, a Log_10_ WBC depletion of 2.88 ± 0.56, and a Log_10_ RBC depletion of 4.19 ± 0.47. A schematic representation of the CTC analytic platform is shown in **Figure 1A**.

**Figure 1:**
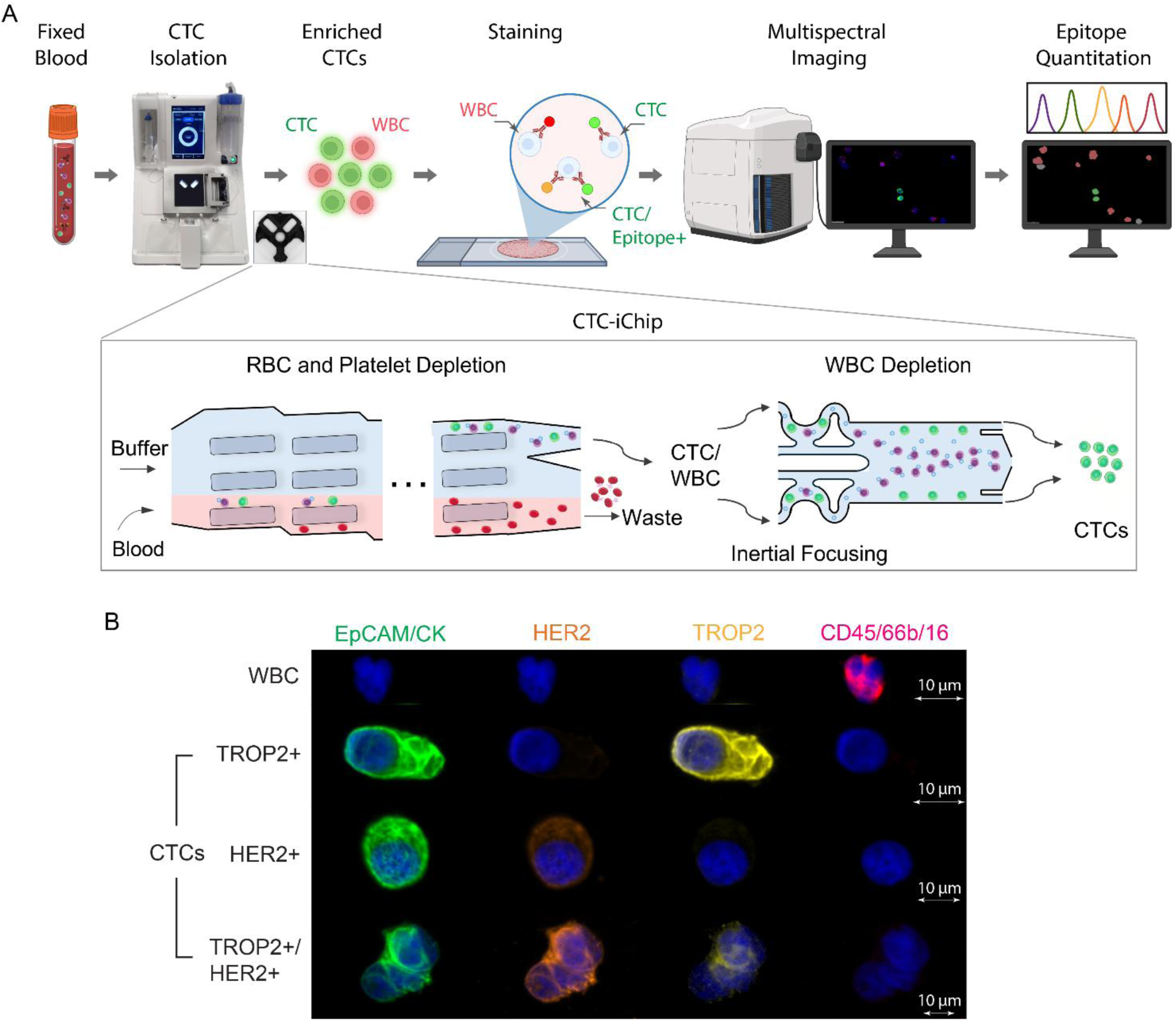
Schema showing the scoring of antibody-drug conjugate (ADC) target expression in Circulating Tumor Cells (CTCs) and representative tumor cells. (A) CTCs from patients with metastatic breast cancer are enriched from 20 mL blood samples collected and fixed using Streck tubes, followed by processing through the microfluidic CTC-iChip for depletion of hematopoietic blood cells (35, 36). Isolated CTCs with residual White Blood Cells (WBCs) are cytospun onto a glass slide, stained with fluorescent antibodies against ADC epitopes of interest, together with shared epithelial/lineage markers (positive identification of CTCs) and WBC markers (negative markers), followed by whole slide imaging (40X) using a multispectral imaging system. For imaging analysis, tumor cells are identified as DAPI+, EpCAM/pan-CK/CK19+, and CD45/66b/16-, and these CTCs are then interrogated for TROP2 and HER2 expression using digital image processing. The inset image shows a schematic diagram of the CTC-iChip system. Red Blood Cells (RBCs), platelets, and plasma are removed using an inertial separation array device, followed by the removal of magnetically tagged WBCs using a magnetic sorter. Nucleated cells are inertially ordered into a single row before flowing into the magnetic sorting channel to provide high-efficiency single-cell sorting, discarding antibody-bound WBCs and collecting unlabeled CTCs. (B) Representative images of CTCs isolated from peripheral blood samples of patients with metastatic breast cancer. CTCs staining for the nuclear marker DAPI (blue), and the general tumor/epithelial lineage markers (EpCAM, pan-CK, and CK19 (Alexa Fluor 488; green)), as well as TROP2 (Alexa Fluor 555; yellow) and HER2 (Alexa Fluor 594; orange) are shown. A WBC staining for CD45, CD66b, and CD16 (Alexa Fluor 647; red) is shown as a negative control.

CTCs are classically defined as cells in circulation that are positive for epithelial markers and negative for leukocyte markers (40). To quantitatively measure the expression of ADC-targeted epitopes across all detectable CTCs, we applied high-resolution (40X) multispectral imaging, taking advantage of sharp emission spectra that allow for scoring multiple epitopes with minimal cross-fluorescence (41). Breast CTCs were identified as intact, nucleated, DAPI-positive cells that also stain positive for one or all of the Epithelial Cell Adhesion Molecule (EpCAM), pan-cytokeratin (CK), or cytokeratin 19 (CK19) markers (all grouped in the Alexa Fluor 488). Cells were simultaneously stained using antibodies against TROP2 (Alexa Fluor 555) and/or HER2 (Alexa Fluor 594) to score for CTCs expressing these ADC targets (**Supplementary Table 1**). Commonly used standard breast cancer cell lines with high, medium, or low expression of TROP2 or HER2 were used to scale the quantification of these markers in CTCs (**Supplementary Figure 2**). Digital image segmentation analysis was used to identify initial CTC images, which were then confirmed manually. Representative images of EpCAM/CK-positive CTCs expressing TROP2 and HER2 are shown in **Figure 1B**.

The clinical characteristics of the 35-patient cohort are shown in **Supplementary Table 2**. Among these, 22 (63%) had hormone receptor-positive (HR+) breast cancer, 11 (31%) had triple-negative (TNBC) disease, and 2 (6%) had evidence of HER2 amplification (HER2+). The mean number of prior therapy regimens was 3.3 (range 0-9). Among all patients studied at the pretreatment baseline, 28 of 35 (80%) cases had ≥3 detectable CTCs within 20mL of whole blood that were of sufficient quality for imaging analysis. Among these 28 cases, the mean number of CTCs per case was 136.0/20mL (6.80 CTCs/mL), with a median of 47 CTCs/20mL (2.35 CTCs/mL) and a range of 4 to 1,481 CTCs/20mL. Of the seven patients with insufficient CTC numbers for analysis at the pretreatment baseline, five had no CTCs (four in the TROP2-ADC cohort and one in the HER2-ADC cohort), and two patients had only two CTCs (HER2-ADC cohort).

### Comparison of TROP2 and HER2 protein expression on CTCs with matched tissue biopsies

Having optimized parameters for CTC quantitation and imaging in MBC, we compared the expression of TROP2 and HER2 epitopes in a separate collection of cases for which patient-matched metastatic tumor biopsies were available. For optimal comparison, we used multispectral fluorescence-based imaging at the single-cell level for both CTCs and matched tumor biopsies, using the same fluorescent-conjugated antibody cocktails. Among all cases selected for this comparison, 30% of metastatic tumor biopsies were of insufficient quality for analysis, while CTCs were too few for analysis in 20% of cases. We were able to analyze both TROP2 and HER2 expression across technically adequate samples for seven matched biopsies and CTCs. In five of these cases, CTC collection was performed within three months of clinically mandated biopsy. Individual CTCs, identified as positive for EpCAM, pan-cytokeratin and CK-19, were scored as null, low, medium, or high for both TROP2 and HER2 proteins, using comparison with established clonal cancer cell lines as controls for staining intensity of these target epitopes (**Supplementary Figures 2 and 3A-C**; see Methods). Given the much larger number of cancer cells within biopsy samples, EpCAM, pan-cytokeratin, and CK-19-positive cells were pooled into null, low, medium, and high quartiles for TROP2 and HER2 expression, with individual biopsies scored for the fraction of cancer cells within each of these expression levels (**Supplementary Figures 3B-C**). For both tumor biopsies and CTCs, the heterogeneity of single-cell expression levels for TROP2 and HER2 was considerable, both within individual cases and across different patients in this cohort, precluding a direct comparison between CTCs and tumor biopsies **(Supplementary Figure 3D-E)**. Both CTCs and tumor biopsies had substantial fractions of single cancer cells ranging from null to high for expression of TROP2 and HER2, without evidence for a clear cut-off applicable across individual patients. Compared with CTCs, tissue biopsies yielded a larger proportion of cells with high expression for both epitopes, with evident discordance in five cases for TROP2 (MGH-022, MGH-039, MGH-040, MGH-042, MGH-043) and one discordant case for HER2 (MGH-040).

Biopsy-based studies have reported a poor correlation between ADC response and expression of either TROP2 (13, 20, 21) or non-amplified HER2 (23, 24) in tumor biopsies. Consistent with the apparent differences between biopsy and CTC scoring, we recently observed a striking discordance in the response of small cell lung cancer (SCLC) to the bispecific antibody tarlatamab targeting the neuroendocrine marker DLL3, with a strong predictive value for DLL3 quantitation on CTCs but not for IHC staining of tumor biopsies (42). We therefore tested quantitative single-cell scoring of CTCs for TROP2 and HER2 epitope expression in MBC patients receiving the respective ADCs.

### Serial CTC enumeration and TROP2 staining in ADC-treated patients

We prospectively studied 13 patients with MBC (**Supplementary Table 2**) treated with a TROP2-targeting ADC as their first antibody-based therapy, analyzing the number of CTCs and their individual cell TROP2 expression levels immediately before initiation of therapy, at three weeks on treatment, and at the time of disease progression. Treated patients were classified as having a poor response if they were maintained on therapy for less than 6 months (four patients; 30.8%) or as having a strong response if they continued therapy beyond that timepoint (nine patients; 69.2%). Four of the strong responders stayed on therapy for over a year (**Figure 2A**). Among all cases, ≥3 CTCs/20mL were detectable at the pretreatment baseline in 9 (69.2%) cases (mean:66 ± 70 CTCs/20mL, median: 34 CTCs/20mL, range: 0 to 217 CTCs/20mL). TROP2 expression within single CTCs was classified as null, low, medium, or high **(Supplementary Figure 2A)**, and cases were scored as TROP2^>50^ if 50% of more CTCs expressed any detectable epitope (i.e., TROP2-low, medium, or high), or as TROP2^<50^ if greater than 50% of CTCs expressed no detectable epitope (i.e., TROP2-null). Consistent with prior results from tumor biopsies (13, 20, 21), the relative expression of TROP2 on CTCs was not correlated with the duration of response to TROP2-ADC. Of the five patients scored as TROP2^>50^, two had a strong response and three had a poor response; of the four patients scored as TROP2^<50^, two had a strong response and two had a poor response (**Figure 2A**). Analysis of a 90% cutoff for TROP2-expressing CTCs similarly did not distinguish strong from poor responders. We cannot exclude a small number of TROP2-expressing cancer cells within a tumor mediating a bystander effect on surrounding cells lacking TROP2 expression.

**Figure 2:**
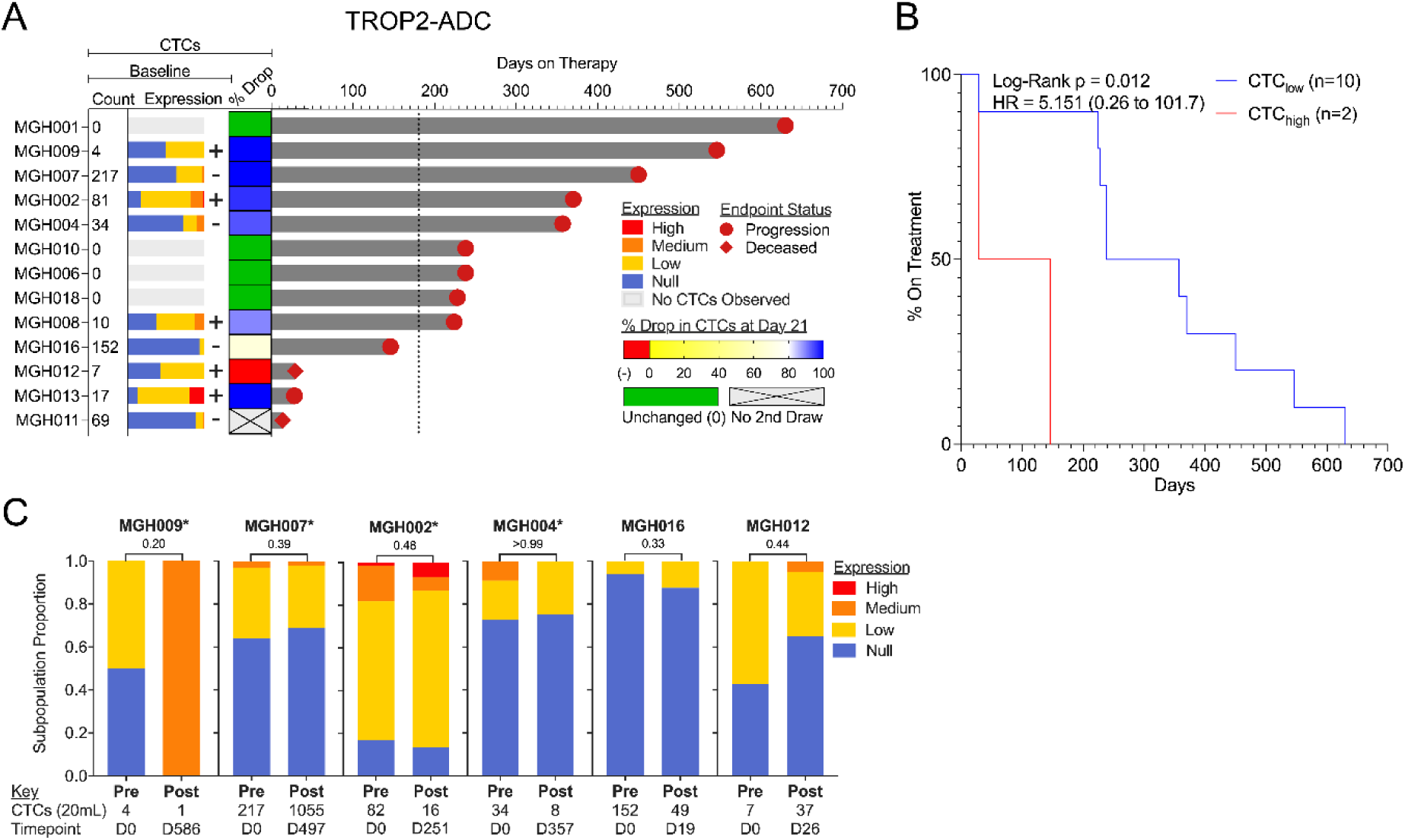
Clinical response and CTC analyses in TROP2-ADC-treated patients. **(A)** Analysis of 13 patients with metastatic breast cancer treated with TROP2-targeting ADCs. From left to right: Number of CTCs isolated from blood (count) at pretreatment baseline, normalized to 20mL; Colored stacked bar graph showing the fraction of CTCs, each expressing undetectable (null), low, medium, and high levels of TROP2 at baseline for each patient (color code shown at right); Scoring of CTC samples as + (>50% of CTCs expressing low, medium or high levels of TROP2) or – (>50% of CTCs expressing no TROP2). Percentage drop in CTC counts at the first post-treatment visit (approximately day 21) for each patient (color code shown at right); Swimmer plot showing the duration of TROP2-ADC therapy (days) as a measure of continued treatment efficacy (Time to Progression), with the reason for treatment discontinuation (progression or deceased) noted. The hashed line represents the six-month distinction between strong and poor responses. **(B)** Kaplan-Meier plot showing time to progression on TROP2-ADC therapy in patients who either have no detectable CTCs or > 80% decline in CTC counts after three weeks of therapy initiation (blue), compared with patients with a lesser decline or an increase in CTC counts at three weeks (red). Log-Rank P-value and hazard ratio (HR) compare ‘>80%’ and ‘<80%’ groups. Evaluable patients had clinical or radiographic assessments and pre- and on-therapy blood draws. (**C**) Persistence of TROP2 expression in CTCs at the time of progression. Bar graphs show the distribution of TROP2 expression in CTCs collected from six patients, comparing pretreatment initiation (left) with matched samples from the same patients at the time of disease progression and at therapy termination. The number of CTCs (normalized to 20mL) and the timepoint of each draw relative to treatment initiation are noted below the plots. Asterisk denotes cases who had a strong initial clinical response to ADC therapy.

While baseline TROP2 epitope expression is not a predictor of ADC response in this cohort, measuring the number of CTCs at the pretreatment baseline and following the first course of therapy may provide a clinically useful measure of early therapeutic benefit, allowing treatment adjustments before the often-delayed radiographic evidence of treatment response. Four of 13 (30.7%) cases had no detectable CTCs (0 CTCs/20mL) at the pretreatment baseline, all of whom remained negative for CTC detection at three weeks on-treatment, and all (100%) went on to have a strong clinical response to TROP2-ADC (**Figure 2A**). Out of six patients who had a >80% decline in CTC numbers at the three-week on-treatment timepoint, five (83.3%%) had a strong response to TROP2-ADC therapy, and one (16.7%) had a poor response. Neither of the two (16.7%) patients with either an on-treatment increase in CTCs or a <80% decline had a strong response to the TROP2-ADC (0%). Together, classifying patients as CTC^Low^ if they have undetectable CTCs or a >80% decline in CTCs after the first treatment dose, nine of ten (90%) CTC^Low^ cases went on to have a strong clinical response and one (10%) had a poor response to TROP2-ADC. In contrast, of two CTC^High^ cases, all (100%) had a poor response (**Figure 2B**). The median duration of response for CTC^Low^ patients was 297 days, compared with 87 days for CTC^High^ cases (Hazard ratio 5.15, p = 0.012). Since only a few patients in our small cohort had the adverse CTC^High^ biomarker, confirmation of these findings in a larger study would be required nominate CTC enumeration for consideration as an early marker of response for TROP2-targeting ADC.

Beyond their ability to predict early response to targeted therapies, the contribution of biomarkers to acquired resistance may also identify critical drivers of drug sensitivity. In six cases, paired CTC specimens were available to compare TROP2 expression at the pre-treatment baseline, versus the time of overt clinical progression. In three of four patients who had a strong clinical response before ultimately experiencing progression on TROP2-ADC and in the two patients with a poor initial response, the single-cell distribution of TROP2 expression on CTCs was unchanged between the drug-sensitive baseline and acquired resistance. In one case (MGH-009) with a prolonged 546-day clinical response, only one CTC, with increased TROP2 expression was detectable at progression (**Figure 2C and Supplementary Figure 4**). Taken together, loss of TROP2-expressing CTCs was not observed in our cohort, coincident with acquired resistance to TROP2-targeting ADCs.

### Serial monitoring of HER2 expression on CTCs in ADC-treated patients

We extended our analysis to an additional 15 patients receiving first-line ADC treatment with the HER2-targeting T-DXd. Among these cases, 13 were classified clinically as having HER2-low tumors (i.e., lacking *HER2* gene amplification, with moderate levels of protein expression), representing the most common indication for T-DXd therapy. *HER2* gene amplification (classical HER2+ breast cancer) denotes high level expression, as well as a biological dependence on HER2 signaling, associated with epitope-dependent response to therapy (43, 44). One patient (MGH-028) had classical HER2+ breast cancer (FISH+ *HER2* gene amplification) and had progressed after first receiving the standard-of-care HER2-targeting antibodies Trastuzumab and Pertuzumab; a second patient’s tumor (MGH-042) lacked *HER2* gene amplification at diagnosis, but a subsequent metastatic tumor biopsy showed heterogeneous foci of FISH-confirmed *HER2* amplification, suggesting the emergence of genetic HER2+ heterogeneity (45). This patient was also previously treated with Trastuzumab. Of the 15 MBC patients, seven (46.7%) were classified as having a strong response to HER2-ADC (remaining on treatment for more than 6 months); this included the two patients with HER2+ (*HER2*-amplified) tumors. Eight patients (53.3%) within this cohort had a poor response (**Figure 3A**). CTCs (≥3 CTCs/20mL) were detectable at the pretreatment baseline in 12 patients (80%) (mean: 216 ± 393 CTCs/20mL, median: 80 CTCs/20mL, range 0 to 1481 CTCs/20mL). HER2 staining was not performed in one of the baseline CTC samples.

**Figure 3:**
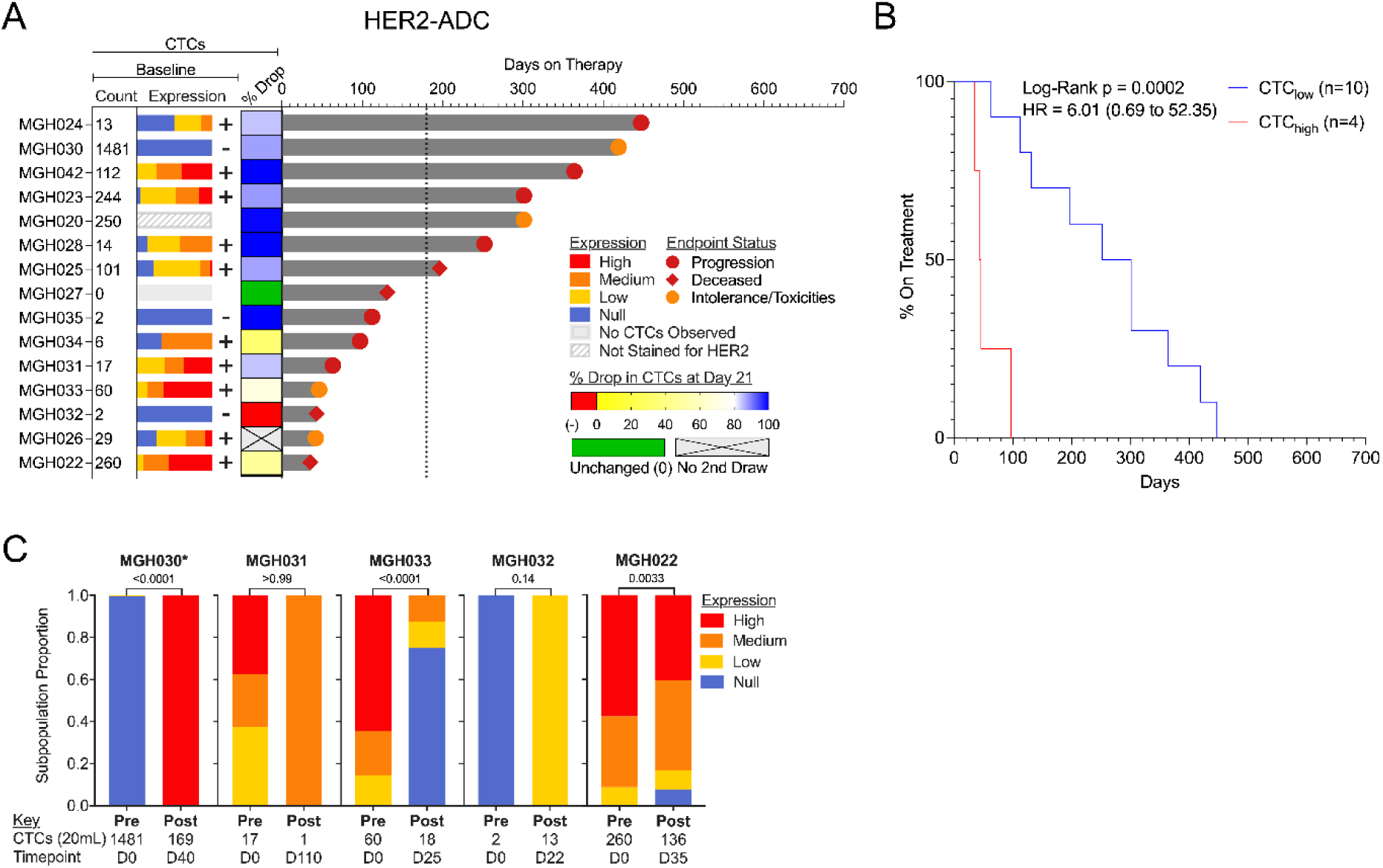
Clinical response and CTC measurements in HER2-ADC-treated patients. **(A)** CTC characteristics and Swimmer Plot analysis of 15 patients with metastatic breast cancer treated with HER2-targeting ADC. From left to right: Number of CTCs isolated from blood (count) at pretreatment baseline, normalized to 20mL; Colored stacked bar graph showing the fraction of CTCs expressing undetectable (null), low, medium, and high levels of HER2 at baseline for each patient (color code shown at right); Scoring of CTC samples as + (>50% of CTCs expressing low, medium or high levels of HER2) or – (>50% of CTCs expressing no HER2). Percentage drop in CTC counts at the first post-treatment visit (approximately day 21) for each patient (color code shown at right). **(B)** Kaplan-Meier plot showing time to progression on HER2-ADC therapy in patients who either have no detectable CTCs or >80% decline in CTC counts after three weeks of therapy initiation (blue), compared with patients with a lesser decline or an increase in CTC counts at three weeks (red). Log-Rank P-value and hazard ratio (HR) compare ‘>80%’ and ‘<80%’ groups. Evaluable patients had clinical or radiographic assessments and pre- and on-therapy blood draws. **(C)** Bar graphs showing the distribution of HER2 expression in CTCs collected from five patients prior to initiation of treatment and at disease progression. CTC number (normalized to 20mL) and time points of each draw relative to treatment initiation are noted below the plots. Asterisk denotes a case that had a strong clinical response to ADC therapy.

At the single cell level, HER2 expression on CTCs was heterogenous, with 10 of 15 (67%) patients having >50% of their single CTCs with detectable (i.e., HER2-low, medium or high) expression (HER2^>50^), three patients (20%) having <50% of their CTCs with detectable expression (HER2^<50^), and one case (6.5%) having 0 CTCs while another case (6.5%) was not stained for HER2. The two patients with *HER2*-amplified tumors had >80% of CTCs expressing HER2 protein, and both had strong responses to HER2-ADC. Across the entire cohort, however, pre-treatment HER2 expression on CTCs was not well correlated with the duration of response to HER2-ADC treatment (**Figure 3A**). Of the ten patients with HER2^>50^ CTCs, five had a strong clinical response and five had a poor response; of the three patients with HER2^<50^ CTCs, one had a strong response and two had a poor response. A 90% cutoff for HER2 expression on CTCs similarly did not yield a correlation with duration of clinical response. Since HER2 immunohistochemistry (IHC) staining on breast tumor biopsies is routinely performed as part of clinical care, we were also able to analyze its predictive value in this cohort: consistent with the CTC results, it showed no correlation with ADC response (**Supplementary Figure 5**).

As with TROP2-targeting ADCs, an early and profound on-treatment decline in CTC numbers after initiation of therapy was highly correlated with response (**Figure 3B**). After only three weeks of HER2-ADC treatment, 10 of 15 (67%) patients had either no CTCs or a >80% drop in CTC burden (CTC^Low^): seven (70%) of these had a strong clinical response, while three (30%) had a poor response. All four cases with <80% drop or increased CTC count at the three-week timepoint (CTC^High^) had poor clinical responses (100%). A patient (MGH-026) was excluded from this analysis because a second blood draw could not be obtained. The median duration of response for CTC^Low^ patients was 276 days, compared with 44 days for CTC^High^ cases (Hazard ratio = 6.01; p < 0.001) (**Figure 3B)**. Thus, as for TROP2-ADC treatment, early on-treatment CTC enumeration may provide a potential predictor of strong clinical response for HER2-targeting ADCs, a finding that will need confirmation in larger clinical studies..

Finally, in five patients (one strong responder and four poor responders) with matched pre-treatment and progression CTC samples following HER2-targeting ADC, we did not observe a consistent loss of epitope expression at the time of acquired resistance. The patient who had a strong initial response to HER2-ADC showed a robust increase in HER2-expressing CTCs at disease progression (MGH-030). Among the four patients who had a poor response to HER2 ADC, two showed no change in HER2-expressing CTC fractions upon acquired drug resistance, one had a decrease in HER2-expressing CTCs, and one acquired CTCs with detectable HER2 expression (**Figure 3C** and **Supplementary Figure 4B**). The two patients with *HER2*-amplified cancers (HER2+) did not have blood samples available for analysis at disease progression. For both TROP2- and HER2-ADC treated cases, high-resolution CTC imaging showed unchanged morphological appearance and epitope staining before treatment initiation and after discontinuation of therapy (**Supplementary Figure 6**).

Taken all together, for both TROP2- and HER2-ADCs, CTC^Low^ measurements at three weeks on-therapy appear predictive of the duration of ADC response. However, the fraction of CTCs expressing different levels of the targeted epitope before treatment initiation is poorly correlated with the duration of clinical response and the acquisition of resistance is not associated with consistent loss of epitope expression.

### Sequential treatment with TROP2- and HER2-targeting ADCs

Patients with MBC who experience progression following initial response to either a TROP2- or HER2-ADC are commonly treated with a second ADC targeting the alternative epitope. We analyzed CTCs from five patients who initially received HER2-ADC and switched to second-line TROP2-ADC at the time of treatment failure, and from five patients receiving second-line HER2-ADC after progression on initial TROP2-ADC (**Figure 4A-B**). In comparison to patients treated with either TROP2- or HER2-ADCs as first-line therapy, nine of the ten patients receiving second-line ADCs had poor responses that fell below the 6-months cut-off (median time to progression: 226 days for any first-line ADC therapy vs 48 days for any second-line therapy) (**Figure 4C**). Among these, seven patients had shorter responses to second-line, compared with first-line ADCs. Of note, patients who had shorter second-line responses had received a mean 12.25 cycles of prior first-line ADC therapy, whereas those who had a modest increase in duration of response to a second-line ADC had only received short courses of first-line treatment: 2 cycles (MGH-005) and 3 cycles (MGH-014) (**Figure 4D**). A remarkable outlier (MGH-029) had only received one cycle of first-line TROP2-ADC, which was discontinued due to toxicity, and she went on to have a prolonged (628-day) response to second-line HER2-ADC. Given the fact that both HER2- and TROP2-ADCs carry similar camptothecin-related drug payloads, it is noteworthy that the few patients who had a favorable response to second-line ADCs were those who had only a short drug exposure during first-line therapy. Within this limited cohort, the generally poor clinical outcome of second-line ADC therapy was independent of the number of CTCs or HER2 expression on CTCs (**Figure 4A, B**).

**Figure 4:**
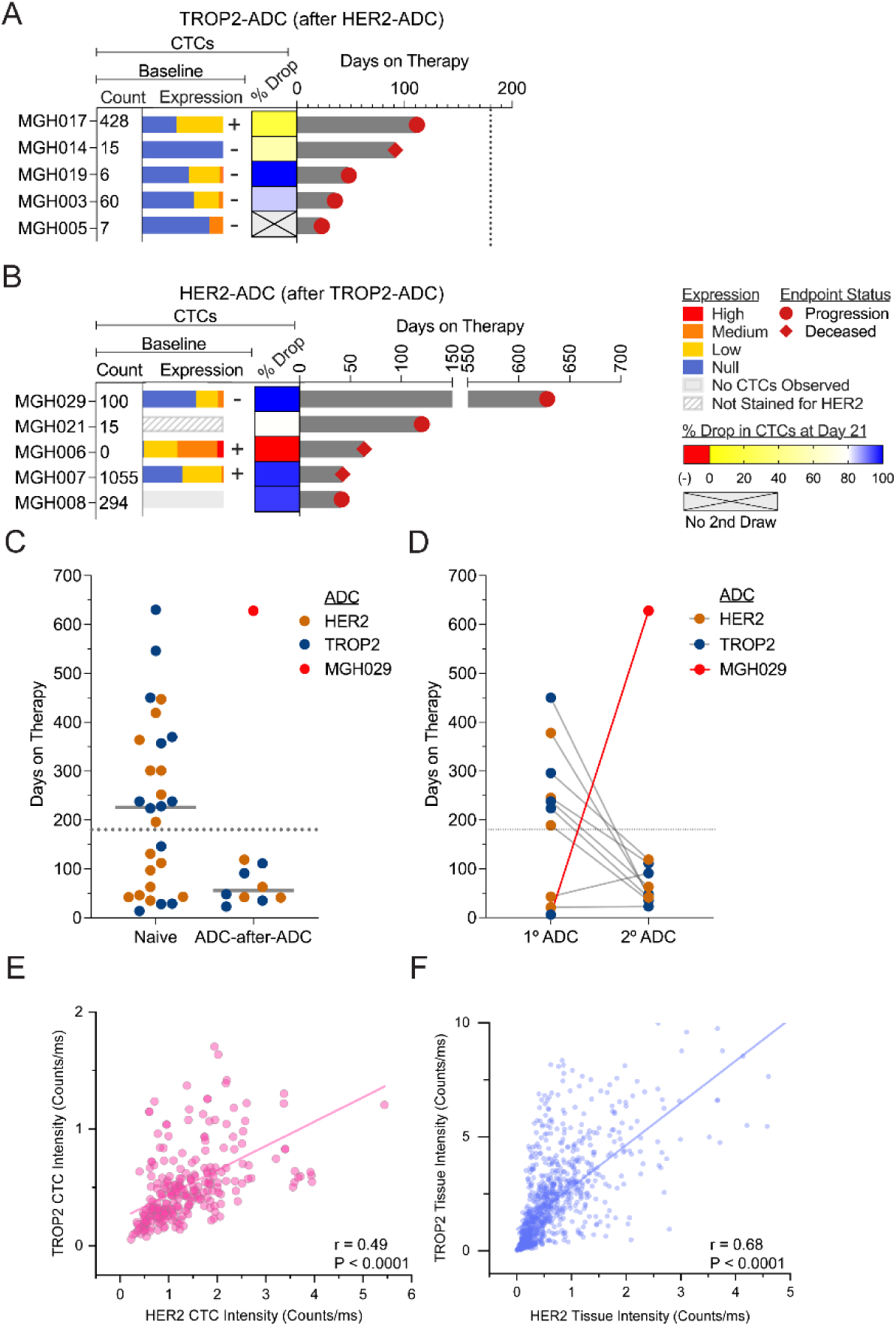
Poor second-line response in patients receiving sequential ADCs. **(A-B)** Analysis of five patients with MBC treated with TROP2-targeting ADC after having progressed on a HER2-targeting ADC (A) and five patients treated with HER2-targeting ADC after having progressed on a TROP2-targeting ADC (B). From left to right: Number of CTCs (count) isolated at pretreatment baseline for the second-line ADC (normalized to 20mL); Colored stacked bar graph showing the fraction of CTCs expressing undetectable (null), low, medium, and high amounts of TROP2 or HER2 (respectively) at baseline for each patient (color codes at right); Scoring of CTC samples as + (>50% of CTCs expressing low, medium or high TROP2 or HER2, respectively) versus – (>50% of CTCs with no detectable expression of TROP2 or HER2, respectively); Percent drop in CTC counts at the first post-treatment visit (approximately day 21) (color code at right); Swimmer plot showing the duration of second-line therapy (days) as a measure of continued treatment efficacy, with the reason for treatment discontinuation (progression or deceased) noted. The vertical dashed line denotes the 6-month cut-off for a strong versus poor response to the second-line ADC. **(C)** Left: Scatter plot showing time to progression (days) for the 10 patients receiving second-line ADC therapy (“ADC-after-ADC”; HER2 followed by TROP2 (n=5) or TROP2 followed by HER2 (n=5)) versus patients receiving first-line ADC therapy (TROP2 (n=13) or HER2 (n=15)). The color code (orange, blue) denotes the ADC used in first-line for the previously untreated (Naïve) cohort and the ADC used in second-line for the “ADC-after-ADC” cohort. The hashed line represents the 6-month cut-off for strong versus poor ADC response, and two solid lines indicate the median time on therapy for each group (first-line: 226 days vs second-line: 48 days; p < 0.001 by Welch’s T-test). Patient MGH029 (red) was excluded from this analysis since first-line TROP2-ADC was terminated after only 6 days (one cycle) due to therapy-related toxicity. **(D)** Plot showing duration on therapy (days) for the 10 patients who received sequential ADC therapies (ADC-after-ADC), comparing their response to first-line versus second-line therapies. **(E-F)** Correlation between TROP2 and HER2 expression within single CTCs (E) and single cancer cells from tumor biopsies (F), pooled from seven ADC naïve cases of metastatic breast cancer, showing the extent of coexpression between these two epitopes within single breast cancer cells.

Given the common clinical scenario of switching between TROP2- and HER2-targeting ADCs, we interrogated breast cancers and CTCs at the single-cell level for coexpression of these tumor-enriched epithelial epitopes. Among the seven cases where matched tumor biopsies and CTCs were stained for both TROP2 and HER2 (**Supplementary Figure 3**), we observed correlations in the expression of these two epitopes within single breast cancer CTCs (r = 0.49, P < 0.0001) and within single tumor cells from metastatic biopsies (r = 0.68, P < 0.0001) **(Figure 4E and F**). Thus, individual breast cancer cells frequently co-express TROP2 and HER2, consistent with their epithelial lineage. Since expression of these epitopes is not generally lost at the time of acquired resistance to either TROP2 or HER2-targeting ADCs **(Figures 2C and 3C)**, switching between these two targeted epitopes while preserving chemically-related drug payloads may not adequately address causes of acquired resistance, leading to the poor clinical outcomes currently observed for second-line ADC therapies.

## DISCUSSION

We describe the clinical application of quantitative single-cell CTC imaging analysis to measure baseline and serial changes in expression of epithelial tumor epitopes targeted by ADCs in patients with breast cancer. The relative ease of blood-based sampling, together with technical improvements in processing and imaging CTCs, enables longitudinal, non-invasive monitoring of protein expression on cancer cells as the tumors initially respond and ultimately progress on ADCs. The emergence of ADCs as a primary therapeutic modality for the treatment of breast cancer brings with it the need to better understand predictive markers of response, both in selecting initial treatment and, following clinical resistance, designing subsequent lines of therapy. For effectively sequencing different ADCs, it is particularly critical to understand the relative importance of substituting different epitope-targeting antibodies versus changing the drug payload conjugated to these antibodies. To our knowledge, this is the first prospective study to define the serial changes in CTC expression of TROP2 and HER2 during the course of ADC therapy. For both TROP2- and HER2-targeting ADCs, we find the greatest potential clinical value for CTC analyses in the rapid measurement of tumor response. Indeed, three weeks after a single administration of either ADC, we find that the persistence of CTCs is highly correlated with a relatively short response, whereas a substantial decline or total absence of CTCs is strongly correlated with a favorable outcome (HR 5.15, p = 0.012 for TROP2 and HR 6.01, p < 0.001 for HER2). Given the poor performance of ADCs administered in the second-line setting, an early marker of tumor response may help in selecting the best first-line ADC treatment.

Whereas early on-treatment CTC numbers, as a measure of tumor burden, are highly predictive of therapeutic response to ADCs, our CTC-based imaging results corroborate previous biopsy studies indicating that pretreatment baseline expression of TROP2 is poorly predictive of ADC response in breast cancer (13, 14, 21). Similarly, for HER2-ADCs, pretreatment epitope staining of CTC is not strongly predictive of ADC response (outside of *HER2*-amplified cases), consistent with biopsy results (6, 7). This confirmation is clinically relevant, since our CTC studies were performed immediately prior to initiation of ADC therapy, excluding potential confounders such as changes in tumor composition over time or selective tumor sampling, both of which may complicate interpretation of metastatic lesion biopsies. Our CTC analysis thus illustrates an emerging paradox in the deployment of TROP2 and HER2-targeting ADCs for breast cancer: ADCs may enable the administration of a highly toxic payload that would not be tolerated without conjugation to an antibody, but the selective distribution and internalization of the drug within cancer cells does not appear to be driven primarily by antibody-dependent targeting of these tumor cells.

Our CTC analysis is most striking for the divergence between the poor correlation of epitope expression and ADC response in breast cancer, compared with the very high concordance of CTC epitope expression and response to a bispecific antibody therapy in small cell lung cancer (SCLC) (42). Using the same CTC isolation, staining, and scoring platforms applied here, we recently tested expression of the neuroendocrine marker DLL3 on CTCs in patients with SCLC treated with the CD3-recruiting, DLL3-targeting bispecific antibody tarlatamab: DLL3-staining of CTCs had 100% specificity and 85% sensitivity for predicting response to the antibody (42). This divergence may provide important insight into two fundamental differences between these antibody-driven cancer therapies. First, the neuro-endocrine DLL3 epitope is highly tumor-selective and virtually absent from normal adult tissues, while both TROP2 and HER2 are epithelial epitopes present on some normal epithelial cells (46–49). Thus, in cases without tumor-specific *HER2* or *TROP2* gene amplification, the more modest differential overexpression of these epitopes on cancer cells may lead to broader tissue pharmacodynamic distribution of antibody. Alternative ADC targets, such as the folic acid receptor present on ovarian cancer cells, appear to have more evidence of tumor cell expression-dependent clinical benefit (50). Second, recruitment of CD3-positive T cells by bispecific antibodies is inherently a localized phenomenon, requiring direct cancer cell contact. In contrast, by virtue of their physical size, ADCs may concentrate at sites of vascular permeability, including tumor deposits, with extracellular proteases releasing the drug payload within the general tumor microenvironment. Such mechanisms may effectively reduce systemic toxicity of ADC payloads but would be less constrained by tumor cell expression of the targeted epitope (21, 23, 51). They also suggest that the chemical composition of the linker tethering the payload to the antibody molecule may prove to be more critical to pharmacodynamic distribution and clinical efficacy than the expression level of the epitope on cancer cells (52–55).

In addition to pretreatment measurements of biomarkers to predict initial clinical response, longitudinal tracking of individual patients through the acquisition of drug resistance can provide unique information as to the drivers of drug susceptibility. In our study, TROP2 and HER2 downregulation were not consistent drivers of acquired resistance to ADCs. Alternative mechanisms of acquired resistance to TROP2-directed ADCs include *TOP1* mutations associated with resistance to the camptothecin drug payload, as well as less well-understood alterations in drug import and metabolism, and membrane trafficking of the epitope (22, 25, 56, 57). For HER2-targeting ADCs, biopsy studies have reported reduced HER2 staining in some cases, albeit with unaltered ADC delivery to the tumor site (6). Chen and coworkers (58) recently reported reduced HER2 staining at treatment progression in half of 102 metastatic tumor biopsies using IHC scoring, and one of 451 patients with a treatment-acquired mutation disrupting HER2 binding. In our CTC analysis using single-cell immunofluorescence staining, one of five patients showed loss of HER2-expressing CTCs at treatment progression; two cases showed an increased fraction of HER2-positive cells; and two had no change in CTC distribution. Taken all together, a reduction in HER2 expression may be evident in a subset of patients who acquire resistance to HER2-ADC.

Our CTC analyses, together with prior biopsy-based studies, suggest that for TROP2- and HER2-targeting ADCs in breast cancer, the drug payload may be more relevant to clinical response or resistance than the targeted epitope itself. Since only a small number of payload molecules are ultimately internalized by each cancer cell, many ADCs make use of TOP1 inhibitors, which are toxic at the minute concentrations achieved through such internalization (19). As a result, currently approved ADCs for breast cancer carry chemically-related payloads, enabling the development of overlapping resistance to diverse ADCs. Such a phenomenon would help explain the poor outcome of current second-line therapeutic strategies switching between TROP2- and HER2-ADCs while retaining a similar drug conjugate, pointing to the importance of developing new ADCs that carry non-cross resistant payloads.

### Limitations of the study

Our conclusions are limited by the small number and heterogeneous group of patients with MBC receiving ADCs as part of their standard-of-care, making this an initial proof-of-principle analysis that will require additional larger clinical studies for confirmation. Our cohort was also focused on heavily pretreated patients, who constitute the majority of patients receiving HER2- and TROP2-ADCs, although these treatments are increasingly being tested earlier in the course of therapy, and our study did not address tumors with *HER2* gene amplification (classical HER2+), in which dependency on HER2 expression is well established as key to therapeutic response. Protein-based liquid biopsies are likely to play an increasing role in assessing the extent of epitope dependence of antibody-mediated therapies, and they may also be combined with DNA sequencing for acquired mutations to understand the diverse mechanisms leading to acquired resistance to ADC therapies.

## Materials and Methods

### Sample Collection – ADC Cohort

Peripheral blood samples from patients were collected into 1 to 2 vacutainers, either into standard EDTA or Streck tubes, according to protocols reviewed and authorized by the Mass General Brigham (MGB) Institutional Review Board (DF-HCC protocol 13-416). Informed consent was obtained from all study participants. All human subjects in this study are females. For the ADC cohort, the inclusion criterion consisted of adult metastatic breast cancer patients starting TROP2-ADC or HER2-ADC therapy. The timepoints of these draws varied between patients due to differing clinical courses but generally coincided with the start of the first two cycles of treatment, radiographic staging, and regular follow-ups. All patients tolerated the venipuncture well. Whole blood donations from healthy donors within our research group were collected in accordance with MGB IRB Protocol # 2009-P-000295. Additional whole blood samples from healthy donors were procured from Research Blood Components, LLC (Brighton, MA).

### CTC-iChip Fabrication and Sample Processing

Upon receipt, the sample volumes were measured, sequentially incubated with antibodies and beads, and then processed through the CTC-iChip. The CTC-iChips used to process samples were fabricated with medical-grade cyclic olefin copolymer (COC) using variotherm injection molding by Stratec Biomedical AG (Germany). Briefly, the devices utilize a pressure-driven flow of whole blood (diluted 1:1 with buffer) containing magnetically labeled WBCs and buffer to enrich CTCs. As the sample passes through the chip, it first flows through 16 inertial separation array devices, which remove the majority of platelets, RBCs, and unbound magnetic beads on the basis of size, allowing CTCs and WBCs to pass. Subsequently, cells are focused into a single-cell stream, and two stages of magnetic depletion channels use magnets outside the chip to separate labeled WBCs from CTCs and unlabeled cells. The final product, concentrated to a fraction of the initial sample volume, is collected while waste fractions are discarded.

Each product containing residual white and red blood cells and any enriched CTCs is kept on ice to preserve cellular morphology. To ensure consistency in downstream work, nucleated cells were enumerated for each sample and split if needed to plate ∼100,000-300,000 cells on each slide. Cells were incubated in 0.5% paraformaldehyde for 10 minutes and cytospun using Epredia™ EZ Megafunnels and a Thermo Shandon Cytospin 4 instrument (2000 rpm, 5 minutes). In lieu of the included Megafunnel slides, TruBond™ 380 slides were used. Following the cytospin, slides were washed once with 1X PBS and air-dried to ensure maximal cell retention. Air-dried slides were stored for 1-2 nights at 4°C until staining.

### WBC depletion antibody preparation

A cocktail of antibodies was utilized to deplete WBCs in samples. These included biotinylated anti-human CD45 (Thermo Fisher Scientific, clone HI30, IgG1, 0.189 µg/million cells), biotinylated anti-human CD16 (BD Biosciences, clone 3G8, IgG1, 0.0189 µg/million cells), and biotinylated anti-human CD66b (Novus Biologicals, clone 80H3, IgG1, 0.0189 µg/million cells). To prepare the cocktail with the necessary concentrations, the three antibodies were diluted into sterile, filtered 1X phosphate-buffered saline (PBS) containing 0.1% bovine serum albumin (BSA) (Sigma-Aldrich #A2058) to improve stability. The cocktail was stored at 4°C and used within 7 days.

### Magnetic beads preparation

1 µm Dynabeads MyOne Streptavidin T1 superparamagnetic beads from Invitrogen were used to magnetically label WBCs with biotinylated antibodies. Before use, beads were washed with three washes of 0.01% Tween-20 (Fisher Scientific BP337-100) in 1X PBS to remove the storage solution, followed by three washes with 0.1% BSA in 1X PBS. Subsequently, beads were stored in the same 0.1% BSA solution at a concentration of 10mg/mL, 10,000,000 beads/µL to minimize nonspecific binding. As with the depletion antibody cocktail, washed beads were stored at 4°C and used within 7 days.

### Buffer preparation

The running buffer for the CTC-iChip, also used for washing and diluting samples and products, was composed of 0.2% Pluronic F-68 (Sigma-Aldrich #K4894) diluted in 1X PBS. Briefly, the polymer was dissolved in PBS until visually dispersed before filtering with a rapid flow 0.22µm filter (Corning #430517). The running buffer was stored at room temperature and used within 2 weeks.

### Staining

For CTC staining, cells were blocked with a solution of 3% BSA and 2% Normal Goat Serum (Abcam #ab7481) in PBS for 1 hour and stained with EpCAM, panCK (CK8/18), and CK19 conjugated to Alexa FluorⓇ 488 (Cell Signaling Technology 5198S, Cell Signaling Technology 4523S, Thermo Fisher Scientific MA5-18158), pooled white blood cell markers CD45, CD16, and CD66b conjugated to Alexa FluorⓇ 647 (BioLegend 302020, BioLegend 304020, BioLegend 305110), TROP2 conjugated to Dylite-550 (Novus Biologicals NBP2-89492R, NBP2-89493R) or TROP2 labeled with Alexa FluorⓇ 555 (abcam ab214488, abcam ab150078), and HER2 labeled with Alexa FluorⓇ 594 (abcam ab11710, abcam ab150160), and DAPI (ThermoFisher Scientific 62248). Staining for all markers was done for 1 hour, using a solution of 0.1% BSA and 1% Tween-20 in PBS as diluent. Conjugated antibodies, DAPI, and any unconjugated primary antibodies were applied first. After two 5-minute washes, secondary antibodies were applied. Final washing was performed sequentially with 0.3% Tween-20 in PBS and 1X PBS before applying mounting media (Thermo-Scientific #P36962), a coverslip, and curing overnight. For the concordance samples, tissue biopsies were preserved in an FFPE block and cut into 5 µm-thick sections. Normal human placenta FFPE slides were used as run controls. Prior to staining, FFPE slides were baked upright in an oven at 65°C for 3 hours. Both concordance CTC and FFPE tissue slides were stained using a Leica Bond RX automated stainer. Prior to blocking, heat-mediated antigen retrieval was performed with Bond ER2 solution (Tris-EDTA buffer pH 9.0) for 40 minutes on the FFPE slides. All incubations were done at room temperature. Slides were coverslipped using ProLong Gold Antifade Mountant with DAPI.

### CTC enumeration

After staining, whole-slide images were acquired using an Akoya Biosciences PhenoImager at 40x magnification (0.25 μm per pixel resolution) in DAPI, AF488, AF555, AF594, and AF647, and autofluorescence channels. Exposure times were guided by prior healthy donors, spiked cells, and kept consistent between samples. Lower exposure scans were used as needed to avoid saturation. These 8-bit images quantified the signal in each channel from 0 to 255. Where necessary, spectral unmixing was applied utilizing the Akoya InForm software. The images were subsequently exported and processed using a HALO image analysis platform. The initial analysis of the entire slide was performed to identify individual cells using nuclear staining and the levels of fluorescence signal in each channel. Adhering to empirically set inclusion and exclusion parameters, customized digital image processing-based segmentation algorithms marked a subset of these cells as possible CTCs requiring further manual verification. These cells were then individually analyzed, considering overall morphology, size, and staining intensity. Following independent analysis by at least two researchers, CTCs were scored. A similar segmentation algorithm was used to mask all CTCs, including nuclei and the entirety of the cytoplasm. The intensity of TROP2 and/or HER2, averaged across this mask and normalized for exposure time, was compared to cut-offs to classify CTCs within specific expression classes. All counts were normalized to 20mL of sample volume.

### CTC Epitope Analysis

As CTCs vary widely in size, morphology, and intensity, a standardized method was developed to organize individual CTCs across a spectrum of HER2 and TROP2 expression into a series of classes, referred to as ‘Null,’ ‘Low,’ ‘Medium,’ and ‘High.’ To ensure these classes were not only consistent from sample to sample but also represented broadly generalizable levels of expression, various cell lines with known expression levels were processed, stained, and analyzed in an identical manner. These included internally established cancer cell lines (BRx-142, breast cancer; Mel-167, melanoma), cultured in RPMI-1640 supplemented with B-27, epidermal growth factor (20ng/mL), fibroblast growth factor (20ng/mL), and antibiotic-antimycotic and incubated in 5% CO_2_ and 4% O_2_, and breast cancer cell lines MDA-MB-231 (RRID:CVCL_0062) and AU565 (RRID:CVCL_1074), sourced from ATCC and cultured as instructed by ATCC. Cut-offs for ‘Low,’ ‘Medium,’ and ‘High’ were based on knowledge of expression levels of these lines, i.e., BRx-142, a CTC line, expresses more TROP2 than most breast cancer lines, while Mel-167 expresses none of the markers and was used as a lower threshold for both TROP2 and HER2. The expression levels in cell lines (BRx142, AU565, MDA-MB-231, and MEL 167) were normalized by exposure and divided into quartiles, categorizing them as low, medium, or high expression. Patient CTCs were then classified according to these thresholds. For tissue specimens, tumor regions of the tissue biopsies were identified by a pathologist. Using a similar quantification approach as was used for CTCs, tumor cells were segmented and phenotyped using HALO software. Cells were identified with DAPI nuclear stain, and the intensity of markers within the nuclear and cytoplasmic compartments for each cell was exported to a data table for statistical analysis. These cut-offs, noted as lines overlaid on violin plots for each respective panel, are shown for reference **(Supplementary Figure 2)**.

### Data availability statement

All images and data will be made available upon request at the Open Science Framework (OSF).

## Acknowledgments

We are grateful to the patients who donated blood to enable this work. This study was supported by NIH grant K25HL169816 (A.M.); NIH grant RO1CA129933 (D.A.H.), NIH grant U01CA214297 (M.T., D.A.H., S.M.), NIH grant R21CA260989 (M.T.), and NIH grant R01CA255602 (M.T., D.A.H., S.M.), the Howard Hughes Medical Institute (D.A.H.), the Breast Cancer Research Foundation (S.M.), and the National Foundation for Cancer Research (D.A.H.).

## Authors’ Contributions

A.M., A.B., D.A.H., S.M., and M.T. developed the concept. A.M., A.B., D.A.H., S.M., Q.E.C., and M.T. designed experiments. A.B., D.A.H., S.M., and M.T. supervised the project. A.M., L.T.N., M.S., Q.E.C., V.R.P., A.D., E.S., K.A.G., K.K., R.B., J.K., K.H.X., and J.W. conducted experiments and analyzed data. R.A., J.F.E., J.W., L.E., C.S.D., S.H., and D.T.T. provided clinical research samples or reagents and gave technical support. A.M., Q.E.C., A.B., R.A., D.A.H., and S.M. wrote the manuscript. A.M., R.A., and Q.E.C. contributed equally to this work.

