## Supplementary File for "Dynamic Monitoring of Antibody Drug Conjugates Targeting TROP2 or HER2 in Breast Cancer using Circulating Tumor Cells"

#### Conflict of Interest Statement

M.T., D.A.H., S.M., and D.T.T. are co-founders of TellBio, a biotechnology company commercializing the CTC-iChip technology. All authors' interests were reviewed and managed by Massachusetts General Hospital and Mass General Brigham in accordance with their conflict-of-interest policies.

**Supplementary Table 1: Antibody vendor information**

| Target | Antibody Name | Vendor | Catalog # | Leica Bond Rx Working Concentration (µg/mL) | Manual Staining Working Concentration (µg/mL) |
|---|---|---|---|---|---|
| EpCAM | EpCAM (VU1D9) Mouse mAb (Alexa Fluor® 488 Conjugate)<br>RRID:AB_10692105 | Cell Signaling Technology | 5198S | 1 | 5 |
| panCK | Pan-Keratin (C11) Mouse mAb (Alexa Fluor® 488 Conjugate)<br>RRID:AB_836889 | Cell Signaling Technology | 4523S | 1 | 5 |
| CK19 | Cytokeratin 19 Monoclonal Antibody (A53-B/A2), Alexa Fluor™ 488<br>RRID:AB_2539532 | ThermoFisher Scientific | MA5-18158 | 1 | 5 |
| CD16 | Alexa Fluor® 647 anti-human CD16 Antibody<br>RRID:AB_492976 | BioLegend | 302020 (100 tests) | 1 | 5 |
| CD45 | Alexa Fluor® 647 anti-human CD45 Antibody<br>RRID:AB_389336 | BioLegend | 304018 (100 tests) | 0.4 | 5 |
| CD66b | Alexa Fluor® 647 anti-human CD66b Antibody<br>RRID:AB_2563171 | BioLegend | 305110 (100 tests) | 2 | 5 |
| Trop2 | TROP-2 Antibody (001) [DyLight 550]<br>RRID:AB_3435918 | Novus Biologicals | NBP2-89492R | Not used | 5 |

| | | | | | |
|---|---|---|---|---|---|
| Trop2 | TROP-2 Antibody (020) [DyLight 550]<br>RRID:AB_3435946 | Novus Biologicals | NBP2-89493R | Not used | 5 |
| Trop2 | Anti-Trop2 antibody [EPR20043]<br>RRID:AB_2811182 | abcam | ab214488 | 5.5 | 5 |
| HER2 | Anti-ErbB2 / HER2 antibody [ICR12] | abcam | ab11710 | 5 | 5 |
| Rabbit IgG | Goat Anti-Rabbit IgG H&L (Alexa Fluor® 555)<br>RRID:AB_2722519 | abcam | ab150078 | 4 | 5 |
| Rat IgG | Goat Anti-Rat IgG H&L (Alexa Fluor® 594)<br>RRID:AB_2756445 | abcam | ab150160 | 4 | 5 |

* Pan-Keratin (C11) Mouse mAb (Alexa Fluor® 488 Conjugate) detects endogenous levels of total keratins 4, 5, 6, 8, 10, 13 and 18. The antibody does not cross-react with other keratins.

### Supplementary Table 2: Patient cohort

| Study ID | Cohort | Lines of Prior Therapy | Target of Prior ADC | Receptor Classification | HER2 IHC | HER2 FISH | Days on Therapy | Baseline CTCs (normalized to 20mL) |
|---|---|---|---|---|---|---|---|---|
| MGH001 | Primary TROP2-ADC | 4 | None | ER+/PR+/HER2- | 2+ | Not Amplified | 630 | 0.0 |
| MGH002 | Primary TROP2-ADC | 3 | None | TNBC | 2+ | Not Amplified | 370 | 81.5 |
| MGH003 | ADC-after-ADC TROP2 | 5 | HER2 | ER+/PR+/HER2- | 2+ | Not Amplified | 35 | 60.0 |
| MGH004 | Primary TROP2-ADC | 4 | None | ER+/PR+/HER2- | 0 | N/A | 357 | 34.4 |
| MGH005 | ADC-after-ADC TROP2 | 3 | HER2 | TNBC | 1+ | N/A | 23 | 6.8 |
| MGH006 | Primary TROP2-ADC | 3 | None | ER+/PR-/HER2- | 0 | N/A | 238 | 0.0 |
| MGH006 | ADC-after-ADC HER2 | 4 | TROP2 | ER+/PR-/HER2- | 0 | N/A | 63 | 293.5 |
| MGH007 | Primary TROP2-ADC | 5 | None | ER+/PR-/HER2- | 2+ | Not Amplified | 450 | 217.0 |
| MGH007 | ADC-after-ADC HER2 | 6 | TROP2 | ER+/PR-/HER2- | 2+ | Not Amplified | 42 | 1055.3 |
| MGH008 | Primary TROP2-ADC | 3 | None | ER+/PR-/HER2- | 0 | N/A | 224 | 10.3 |
| MGH008 | ADC-after-ADC HER2 | 4 | TROP2 | ER+/PR-/HER2- | 0 | N/A | 41 | 0.0 |
| MGH009 | Primary TROP2-ADC | 0 | None | TNBC | 1+ | N/A | 546 | 4.0 |
| MGH010 | Primary TROP2-ADC | 1 | None | TNBC | 1+ | N/A | 238 | 0.0 |
| MGH011 | Primary TROP2-ADC | 6 | None | ER+/PR-/HER2- | 0 | N/A | 14 | 68.8 |
| MGH012 | Primary TROP2-ADC | 2 | None | TNBC | 0 | N/A | 29 | 6.8 |
| MGH013 | Primary TROP2-ADC | 0 | None | TNBC | 0 | N/A | 28 | 17.2 |
| MGH014 | ADC-after-ADC TROP2 | 3 | HER2 | ER+/PR-/HER2- | 1+ | N/A | 91 | 5.7 |
| MGH016 | Primary TROP2-ADC | 0 | None | TNBC | 0 | N/A | 146 | 152.3 |
| MGH017 | ADC-after-ADC TROP2 | 9 | HER2 | ER+/PR+/ HER2- | 1+ | N/A | 111 | 14.6 |

| | | | | | | | | |
|---|---|---|---|---|---|---|---|---|
| MGH018 | Primary TROP2-ADC | 5 | None | ER+/PR+/HER2- | 0 | N/A | 228 | 0.0 |
| MGH019 | ADC-after-ADC TROP2 | 3 | HER2 | ER+/PR+/HER2- | 2+ | Not Amplified | 48 | 428.0 |
| MGH020 | Primary HER2-ADC | 1 | None | ER+/PR+/HER2- | 1+ | N/A | 301 | 250.0 |
| MGH021 | ADC-after-ADC HER2 | 7 | TROP2 | ER+/PR-/HER2- | 0 | N/A | 119 | 15.0 |
| MGH022 | Primary HER2-ADC | 2 | None | ER+/PR+/HER2- | 2+ | Not Amplified | 35 | 259.5 |
| MGH023 | Primary HER2-ADC | 3 | None | ER+/PR-/HER2- | 1+ | N/A | 301 | 244.5 |
| MGH024 | Primary HER2-ADC | 5 | None | ER+/PR-/HER2- | 2+ | Not Amplified | 447 | 13.2 |
| MGH025 | Primary HER2-ADC | 2 | None | ER+/PR-/HER2- | 2+ | Not Amplified | 196 | 101.3 |
| MGH026 | Primary HER2-ADC | 2 | None | ER+/PR-/HER2- | 1+ | N/A | 42 | 28.5 |
| MGH027 | Primary HER2-ADC | 0 | None | TNBC | 1+ | N/A | 131 | 0.0 |
| MGH028 | Primary HER2-ADC | 2 | None | ER-/PR-/HER2+ | 2+ | Amplified | 252 | 14.3 |
| MGH029 | ADC-after-ADC HER2 | 4 | TROP2 | TNBC | 1+ | N/A | 628 | 99.5 |
| MGH030 | Primary HER2-ADC | 0 | None | ER+/PR-/HER2- | 2+ | Not Amplified | 419 | 1480.5 |
| MGH031 | Primary HER2-ADC | 6 | None | ER+/PR+/HER2- | 2+ | Not Amplified | 63 | 16.9 |
| MGH032 | Primary HER2-ADC | 1 | None | TNBC | 2+ | Not Amplified | 43 | 2.2 |
| MGH033 | Primary HER2-ADC | 7 | None | ER+/PR+/HER2- | 1+ | N/A | 46 | 59.5 |
| MGH034 | Primary HER2-ADC | 6 | None | ER+/PR-/HER2- | 0 | N/A | 97 | 6.5 |
| MGH035 | Primary HER2-ADC | 2 | None | TNBC | 1+ | N/A | 112 | 2.2 |
| MGH042 | Primary HER2-ADC | 2 | None | ER-/PR-/HER2+ | 2+ | Amplified | 364 | 111.9 |

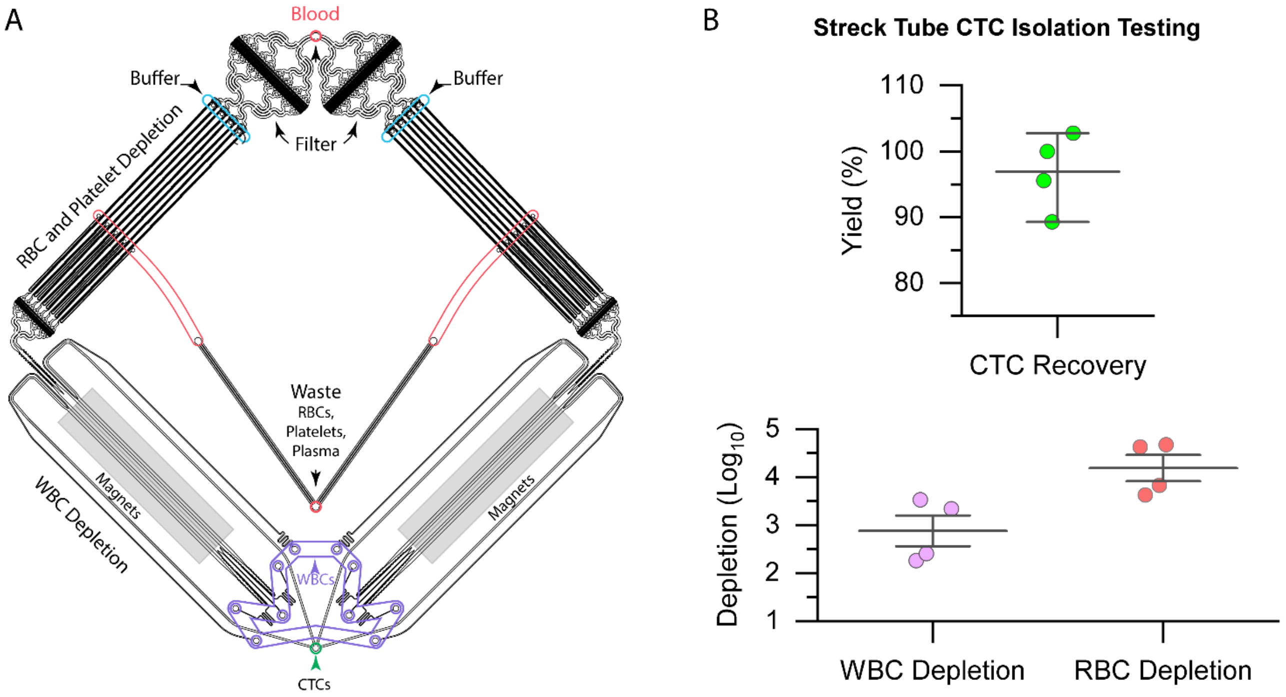


**Supplementary Figure 1**: **CTC-iChip and cell recovery rates (A)** Schematics of the microfluidic CTC-iChip configuration. **(B)** CTC recovery rates and enrichment levels of fluorescence-tagged CTCs from *in vitro* cultures, spiked into blood from healthy donors, fixed using commercial Streck tubes and stored for 24 hrs before processing through the CTC-iChip, showing 96.9 ± 5.1% CTC recovery, 2.88 ± 0.56 ($\log_{10}$) WBC depletion, and 4.19 ± 0.47 ($\log_{10}$) RBC depletion.

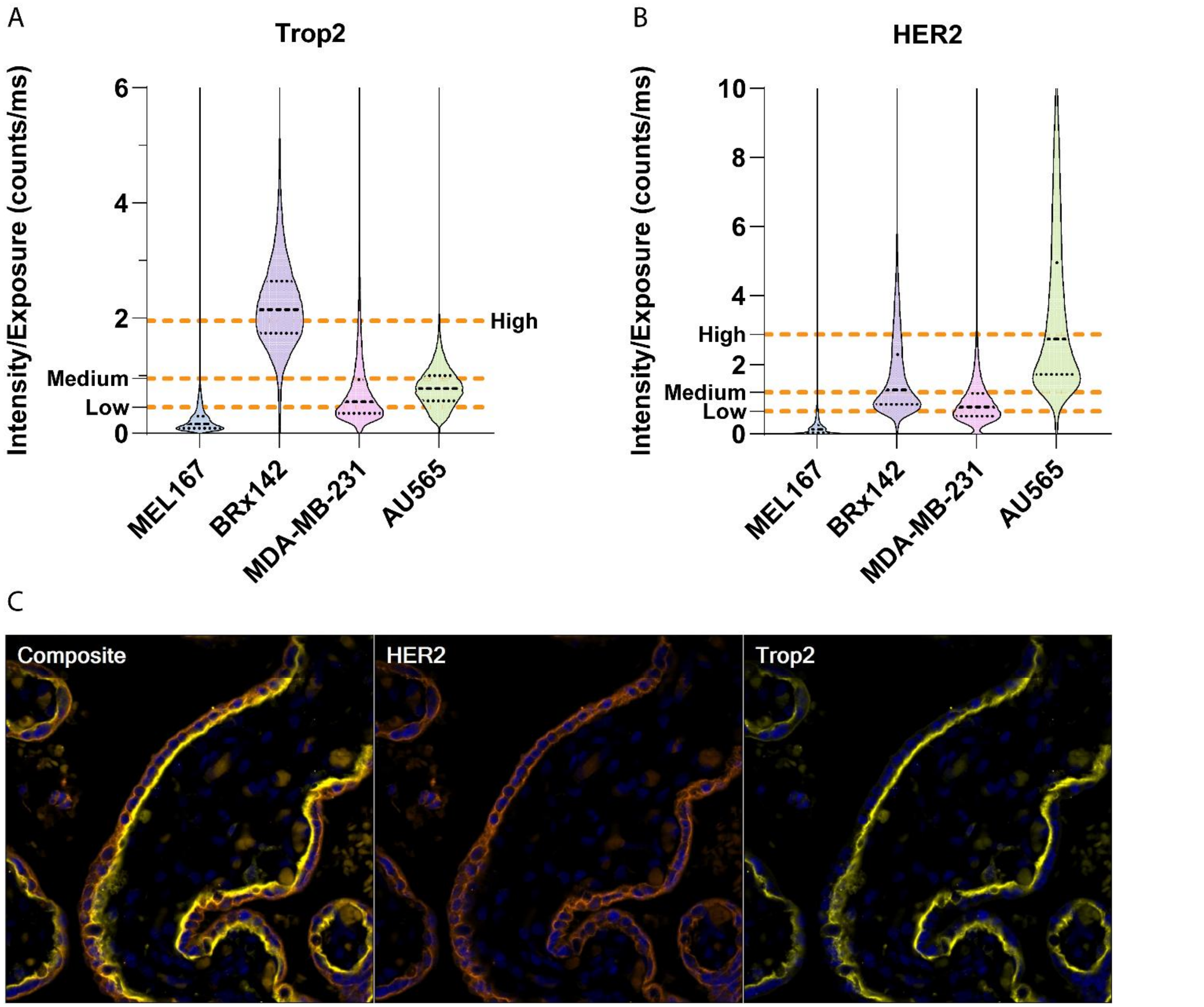


**Supplementary Figure 2**: **TROP2 and HER2 immunofluorescence staining controls (A-B**) Staining intensity of TROP2 (A) and HER2 (B) proteins in the control cell lines shown, used to establish high, medium, and low expression thresholds (dashed lines) applied to scoring CTCs. **(C)** Validation of TROP2 and HER2 antibodies using placental tissues as positive controls.

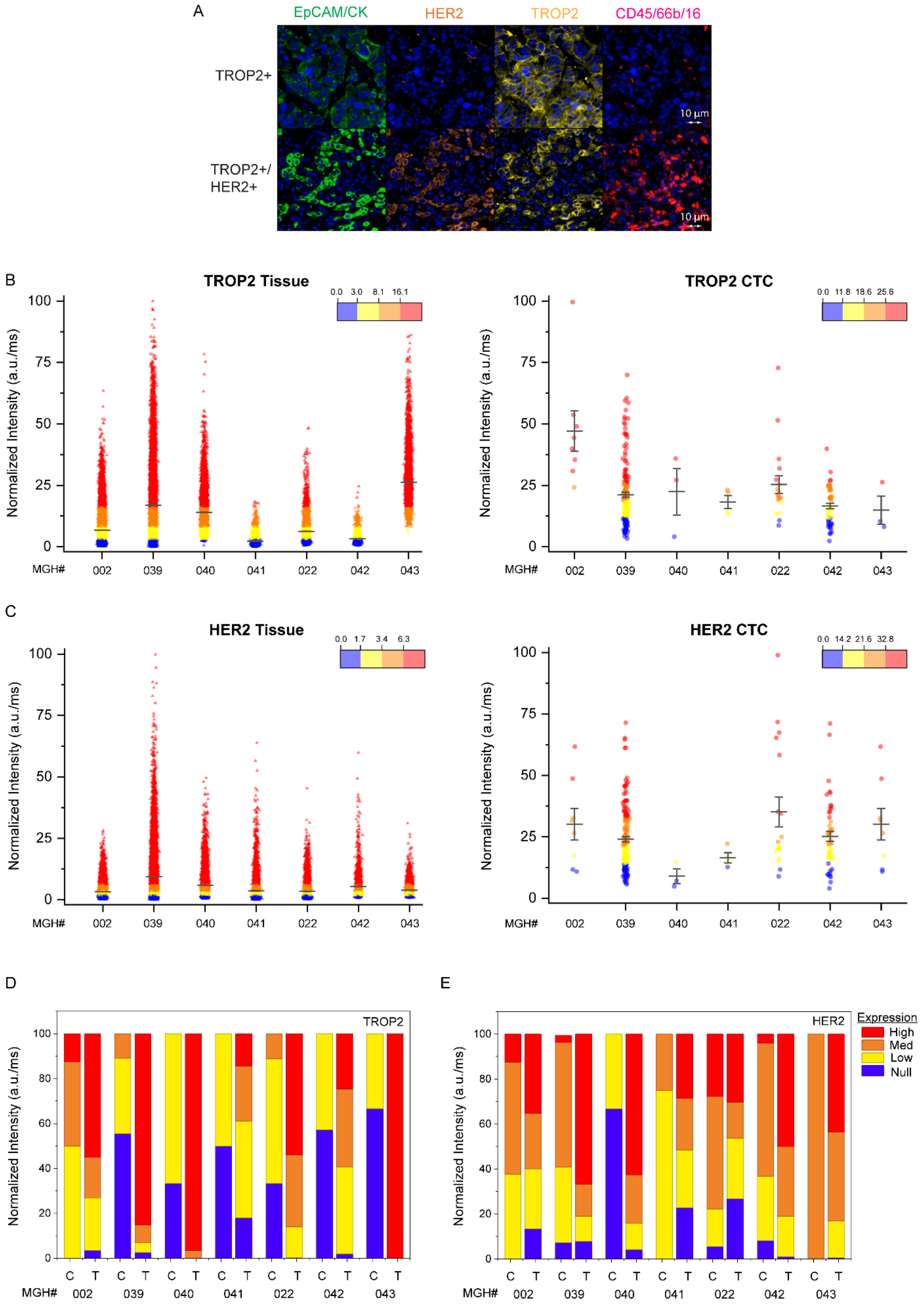
A
EpCAM/CK
HER2
TROP2
CD45/66b/16
TROP2+
TROP2+/ HER2+
10 µm
B
TROP2 Tissue
TROP2 CTC
0.0 3.0 8.1 16.1
0.0 11.8 18.6 25.6
Normalized Intensity (a.u./ms)
MGH#
002 039 040 041 022 042 043
C
HER2 Tissue
HER2 CTC
0.0 1.7 3.4 6.3
0.0 14.2 21.6 32.8
D
TROP2
E
HER2
Expression
High
Med
Low
Null
C T

**Supplementary Figure 3**: **Comparison of TROP2 and HER2 expression in CTCs and matched tissue biopsies. (A)** Immunofluorescence images of a metastatic tumor biopsy stained using the nuclear marker DAPI (blue), and the general tumor/epithelial lineage markers (EpCAM, pan-CK, and CK19 (Alexa Fluor 488; green)), as well as TROP2 (Alexa Fluor 555; yellow) and HER2 (Alexa Fluor 594; orange), and leukocyte markers (CD45, CD66b, and CD16 (Alexa Fluor 647; red)). The upper tumor was positive for TROP2 expression only, while the lower tumor was positive for both TROP2 and HER2 expression. **(B)** Single-cell TROP2 expression in metastatic tumor biopsies (tissue) and in CTCs for seven cases in which such matched paired samples were available. For both CTCs and biopsies, single cells are shown as blue (null expression), yellow (low expression), orange (medium expression), or red (high) TROP2 expression, using the quartile classifications shown in Supplementary Figure 2A. The total number of single cells scored are as follows: MGH002: (biopsy: 24,356; CTCs: 8), 039: (biopsy: 14,720; CTCs: 166), 040: (biopsy: 11,704; CTCs: 3), 041: (biopsy: 5,637; CTCs: 4), 022: (biopsy: 5,061; CTCs; 18), 042: (biopsy: 2,163; CTCs: 49), and 043: (biopsy: 5,617; CTCs: 8). **(C)** Analogous comparison of single-cell HER2 expression in tumor biopsies (tissue) and their paired CTCs, as described above. CTCs and tumor cells were classified as null, low, medium, or high in HER2 expression, using the quartile divisions shown in Supplementary Figure 2B. **(D)** Individual comparisons for each of the seven paired clinical samples, showing the fraction and intensity of TROP2 staining for CTCs (C) versus single cells within tumor biopsies (T). **(E)** Analogous comparison of the same seven paired clinical samples for HER2 expression, comparing CTCs and matched biopsy specimens.

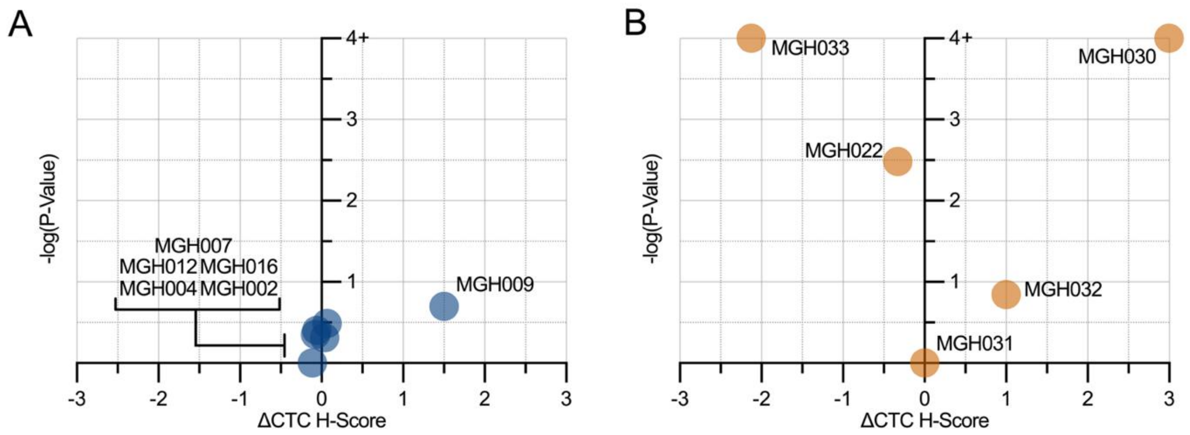


**Supplementary Figure 4: Variation in CTC expression patterns at pretreatment baseline versus clinical progression. (A)** Plot showing both the change in TROP2 CTC H-score from baseline to progression as well as the log(P-value) (Fisher's exact test) for each of six patients on TROP2-ADC, quantifying the significance of changes in epitope expression between matched pretreatment and ADC-resistant CTCs. H-score is defined as 1×proportion of 'Low' cells + 2×proportion of 'Medium' cells + 3×proportion of 'High' cells. Only one out of five patients (MGH009) demonstrated a meaningful (>1) difference in H-score between draws (increased epitope expression at progression). **(B)** Plot showing the change in HER2 CTC H-score from baseline to progression and the log(P-value) (Fisher's exact test) for each of five patients on HER2-ADC. One patient showed a statistically significant decline in HER2 expression, and one patient showed increased HER2 at progression, with the remaining three patients showing no change in CTC expression.

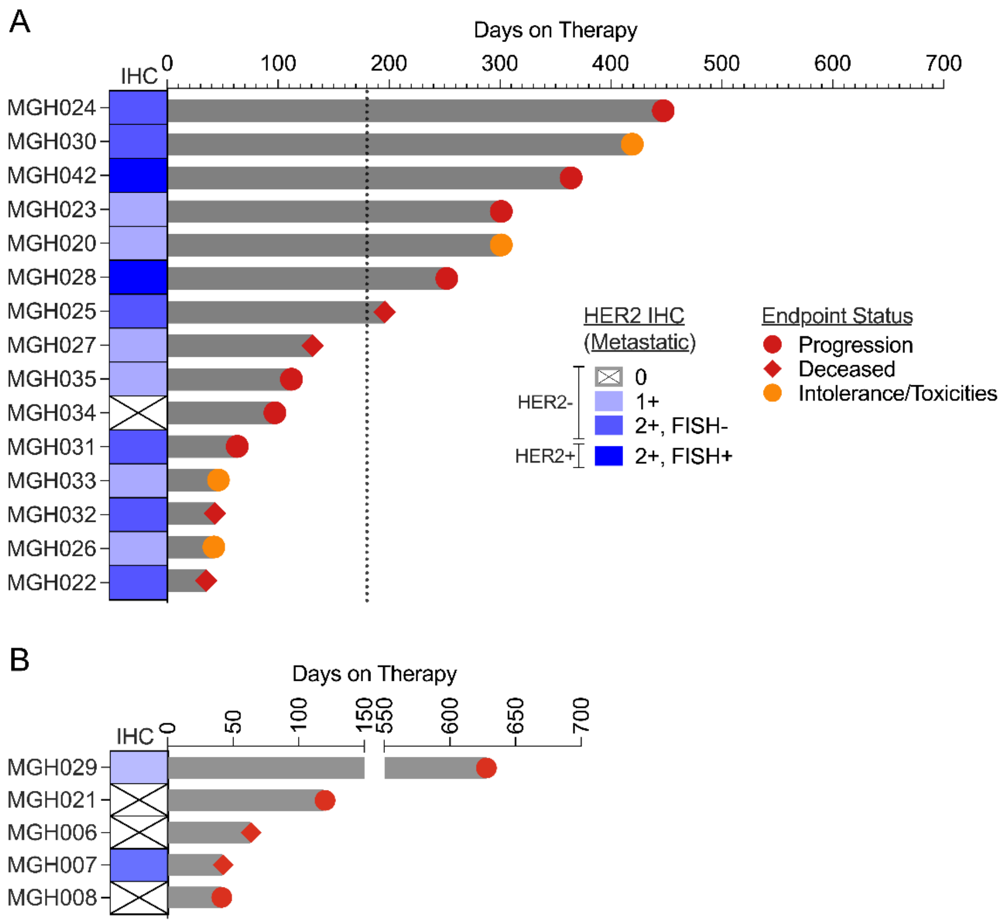


**Supplementary Figure 5 Swimmer plot for tumor-based HER2 IHC staining (A)** Duration of clinical response (time on therapy) for patients treated with HER2-ADC, as a function of routine clinical staining for HER2 using immunohistochemistry (IHC) and *HER2* gene amplification measurements (FISH). **(B)** Duration of clinical response in patients who had previously received TROP2-ADC prior to treatment with HER2-ADC (ADC-after-ADC cohort), as a function of tumor HER2 staining (clinical IHC staining with FISH assay). Patient MGH029 had only previously received one dose of TROP2-ADC, which was discontinued after six days for toxicity.

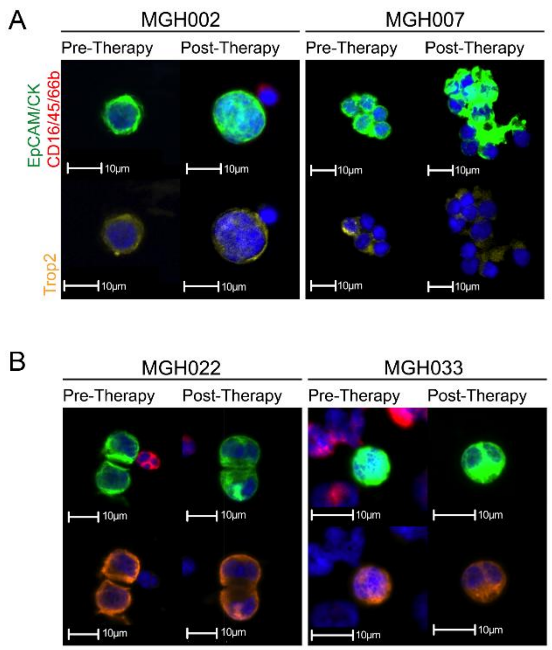


**Supplementary Figure 6**: **Pattern of epitope staining at clinical progression** (A) Representative images of TROP2 expression in patient-matched CTCs from the time of pre-therapy baseline versus clinical progression, demonstrating unchanged cytological features and epitope staining patterns. (B) Representative images showing similar HER2 expression and CTC morphology in similarly paired pre-therapy and progression samples.
